# Cryptic survival and an unexpected recovery of the long-tailed mayfly *Palingenia longicauda* (Olivier, 1791) (Ephemeroptera: Palingeniidae) in Southeastern Europe

**DOI:** 10.1101/2021.04.13.439678

**Authors:** Avar L. Dénes, Romina Vaida, Emerencia Szabó, Alexander V. Martynov, Éva Váncsa, Beáta Ujvárosi, L. Keresztes

## Abstract

1. Once widespread in the large European rivers, *Palingenia longicauda* underwent a drastic range contraction as a result of the intense pollution and hydromorphological interventions of the 19^th^ and 20^th^ centuries. For the last decades it was considered to be restricted only to the Tisa River and its tributaries, and to the Rába River, but new reports indicated its presence in the Danube River in Hungary, in the Danube Delta in Romania and Ukraine, and in the Prut River in the Republic of Moldova.
2. The objective of this study is to analyze the phylogeographic pattern between the two main eco-regions (Pannon and Pontic) of the species distribution, based on the combined mitochondrial COI (472 bp) and 16S (464 bp) sequences generated for individuals collected in Romania and Ukraine, and from publicly available ones, representing the Tisa catchment populations.
3. The presence of viable populations in the Danube Delta and on the Prut River in Romania is confirmed, and additional presence on the Mure□ and Bega rivers from Romania, and on the Styr and Horyn’ rivers in Northern Ukraine is shown.
4. The phylogeographic results indicate that the presence of the analyzed populations are not the result of recent founding events from the Pannon region, confirming the survival and expansion of cryptic local lineages.
5. The recent recovery of the species may be related to the improvement of water quality as a result of the implementation of the EU Water Framework Directive and the EU Floods Directive after 2000.

## Introduction

Freshwater ecosystems are vital resources for people and their livelihoods, having important contributions to the long term management of the environment and communities in a complementary framework (Darwall *et al*., 2011; Voulvoulis *et al*., 2017; Kuntke *et al*., 2020). These ecosystems, especially rivers have major roles in the evolutionary history of an important number of species, as they can act as corridors for riverine organisms, promoting dispersal and diversification (Dijkstra *et al*., 2014). However, freshwater biodiversity is declining at a higher rate than the terrestrial and marine diversity (WWF, 2020; Albert *et al*., 2021), due to changes in land-use (exploitation, habitat degradation, eutrophication, urbanization), flow regulations (channelization, building of dams), or introduction of invasive species (Hein *et al*., 2016; Leese *et al*., 2016; Carrizo *et al*., 2017). Habitat loss or degradation is the most relevant risk factor, affecting 80% of threatened freshwater species (Collen *et al*., 2014). The range loss and fragmentation often results in small isolated populations, leading in time to loss of genetic diversity and to increased extinction risk, by reducing the potential of populations to adapt to possible future challenges like pollution, diseases and climate change (Alexander *et al*., 2011; Werth *et al*., 2014; Pavlova *et al*., 2017; Coleman *et al*., 2018; Dupuis *et al*., 2020).

A well-known example of such dramatic range loss in Central Europe is that of the long-tailed mayfly *Palingenia longicauda* (Olivier, 1791) (Ephemeroptera: Palingeniidae); an iconic species for conservation of pristine riverine ecosystems, and probably the best-known mayfly in Europe, thanks to its huge body size (32–40 mm, up to 100 mm with cerci, and with forewings of 25–37 mm in length) and to the well synchronized mass swarming in mid and late June (Kriska *et al*., 2007; Málnás *et al*., 2011). The life cycle of this species lasts for three years. The larvae prefer steep clay banks, making horizontal U-shaped borrows (Figure 1), and are highly sensitive to organic pollution and riverbank regulations, which lead to the rapid disappearance of the species from the majority of the highly polluted and regularized large rivers of Europe (Russev, 1987). Once widespread and well-known from the lower and middle courses of large and medium-sized rivers all over Europe, the species underwent drastic range contraction that coincided with the intense pollution and hydromorphological interventions that started in the 19th century. To the second half of the 20^th^ century *P. longicauda* was considered extinct in most of its historic range (Russev, 1987; Soldán *et al*., 2009; Bauernfeind & Soldán, 2012). As a result of this near extinction, the species became the most critically endangered mayfly species of Europe, therefore it was included in Appendix II of the Convention on the Conservation of European Wildlife and Natural Habitats (Bern Convention), the Carpathian List of Endangered Species (Witkowski *et al*., 2003) and the Red data book of Ukraine (Akimov, 2009). Conservation efforts include the protection of the species in its known habitats, and an attempt to reintroduce the species on the Lippe River, Germany, which unfortunately was unsuccessful (Tittizer *et al*., 2008; Jourdan *et al*., 2018). The real natural phenomenon of mass swarming of the adults (Figure 1) got important social interest in the past (fishermen used them for bait), as well as in the present, when digitalized information networks, including different social media platforms, document the presence of the species in some remote area, like the Danube Delta or the Prut River in Romania.

**Figure 1.**
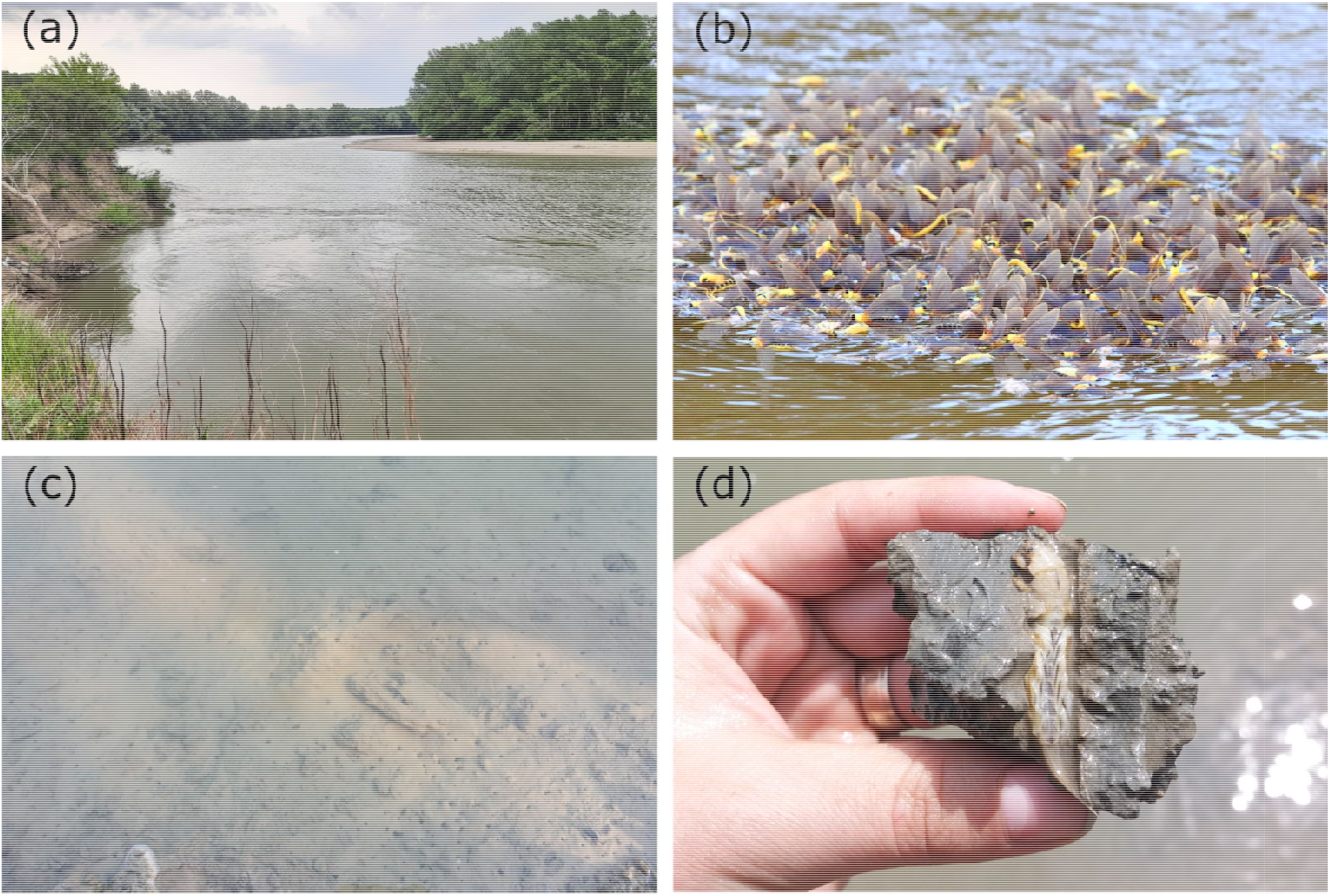
(a) Typical *P. longicauda* habitat with steep clay banks (Mure□ River, Nădlac, Arad county, photo: Vaida R.); (b) group of individuals during the mass swarming (Danube Delta, Maliuc, Tulcea county, photo: Petrescu D.); (c) the openings of the horizontal U-shaped borrows made by the larvae (Prut River, Ia□i county, photo: Vaida R.); (d) larva in the burrow (Mure□ River, Nădlac, Arad county, photo: Vaida R.).

*Palingenia longicauda* is considered a Pontic biogeographical element (Haybach, 1998), which means that the Black Sea region probably had an important role in the species survival. This region is traditionally considered one of the most important refugium for a large variety of organisms during the Pleistocene glaciations (Bănărescu, 1991; Hewitt, 1999). These species dispersed northwards from here, recolonizing Central Europe by following the Danube River and the Danube Basin (Varga, 2010; Bauernfeind and Soldán, 2012). Similarly, extensive phylogeographic studies have shown that the middle-lower Danube catchment within the Pannon region was also an important refugium area for several aquatic biota (Babic *et al*., 2005; Schmitt & Varga, 2012; Vörös *et al*., 2016), including *P. longicauda* (Bálint *et al*., 2012).

For the last decades *P. longicauda* was considered to be restricted only to the Tisa (or Tisza, Tysa) River and the lower range of its tributaries, and to the Rába River (Andrikovics *et al*., 1992; Kovács *et al*., 2001). Bálint *et al*. (2012) published a comprehensive study that included 245 extant specimens from the Tisa and Rába rivers, and their tributaries to assess the loss of genetic diversity caused by the large-scale range loss. They hypothesized that the presence of the species in the Rába River could be the result of recent range expansion. Their results based on a 936-bp long sequence matrix of the combined mitochondrial COI (472 bp) and 16S (464 bp) sequences showed an unexpected high genetic diversity for both rivers, and a significant genetic differentiation between the Tisa (228 specimens) and Rába (17 specimens) rivers. They further show that historic gene flow may have existed between the two rivers, probably before the last glacial maximum (LGM), but that there is no evidence of current connection between them. The authors concluded that the species probably survived the LGM in two middle-Danubian refugia, with a post-LGM introgression event from the lower-middle Danube drainage into populations upstream, suggesting the possibility that the species persisted in the Rába River in small undetected populations.

In recent years several new reports of *P. longicauda* were published showing the presence of the species in the Danube River in Hungary (Málnás *et al*., 2016), in the Danube Delta in Romania (Soldán *et al*., 2009; Bulánková, Beracko and Derka, 2013; Pavel *et al*., 2019) and Ukraine (Afanasyev *et al*., 2020), in the Prut River in the Republic of Moldova (Munjiu, 2018), and in Styr and Horyn’ rivers (Pripyat River basin) in Ukraine (Martynov, 2018 – as *Palingenia fuliginosa* (Georgi, 1802), misidentification). *Palingenia fuliginosa* from Pripyat River basin was recorded by Martynov (2018) based on a morphological study of larvae and subimagoes only. DNA material of specimens mentioned by Martynov (2018) was originally planned to be used as outgroup in the current study, but investigation of their *mt*COI gen and 16S ribosomal rRNA showed that all analyzed specimens belong to *P. longicauda*. This misidentification was also later confirmed by investigation of male imago genitalia.

In the context of these new information, we focused on discovering and sampling *P. longicauda* populations from the major rivers of Romania in order to assess the molecular genetic diversity and the phylogeographic pattern of the species by expanding the molecular framework of Bálint *et al*. (2012) to the whole known distribution of the species.

The objective of this comparative population genetic study is to analyze the phylogeographic pattern between the two main regions of the species distribution, the Pannon and the Pontic regions. The present analyses will therefore focus on two alternative hypotheses:

1. Only a single major LGM refugium of *P. longicauda*, located in the middle sector of the Tisa River (support by literature data, Bálint *et al*., 2012) contribute to the long term preservation of this species in Europe. The sporadic presence of the species in rivers connected to the lower sector of the Danube (including the Danube Delta), is the results of recent colonization events of few individuals from the leading edge of the species.
2. In contrast, the massive presence of the species in the Danube Delta and Prut River, documented in the last years (Soldán *et al*., 2009; Bulánková *et al*., 2013; Martynov, 2018; Munjiu, 2018; Pavel *et al*., 2019; Afanasyev *et al*., 2020), represents overlooked or cryptic populations, suggesting a recovery of some autochthonous populations, and establishing a good ecological status of the aquatic ecosystems in the Danube Delta area and its affluents, because of the implementation of the EU Water Framework Directive after 2000.

## Materials and methods

### Sampling and DNA sequencing

Individuals were collected from 14 locations corresponding to 4 rivers (Danube Delta, Mure□, Bega and Prut) from Romania, and 4 locations corresponding to 2 rivers, Styr and Horyn’ from Ukraine, both tributaries of Pripyat River (Table1 and Figure 2). Genomic DNA was extracted from 196 specimens preserved in 97% ethanol, using a Bioline ISOLATE II Genomic DNA Kit. To be able to integrate the sequence data generated by the previous study (Bálint *et al*., 2012), a 471 base pairs (bp) section of the *mt*COI gen and a 464 bp fragment of the 16S ribosomal rRNA was amplified with the Jerry (Simons *et al*., 1994) – S20 (Pauls *et al*., 2006), and 16Sar (Simons *et al*., 1994) – 16SB2 (Monaghan *et al*., 2007) primer pairs. PCR reactions were performed in a 25 µl reaction volume, at an annealing temperature of 40 °C for the *mt*COI and 56 °C for the 16S fragments, and were sequenced by Macrogen Europe. Sequences were verified at the NCBI website using a Basic Local Alignment Search Tool (Johnson *et al*., 2008) and were deposited in GenBank (accession numbers, *mt*COI: MW716042 – MW716237; 16S: MW717693 – MW717888). Consensus sequences were aligned manually using BioEdit 7.2.5 (Hall, 1999).

**Table 1.**
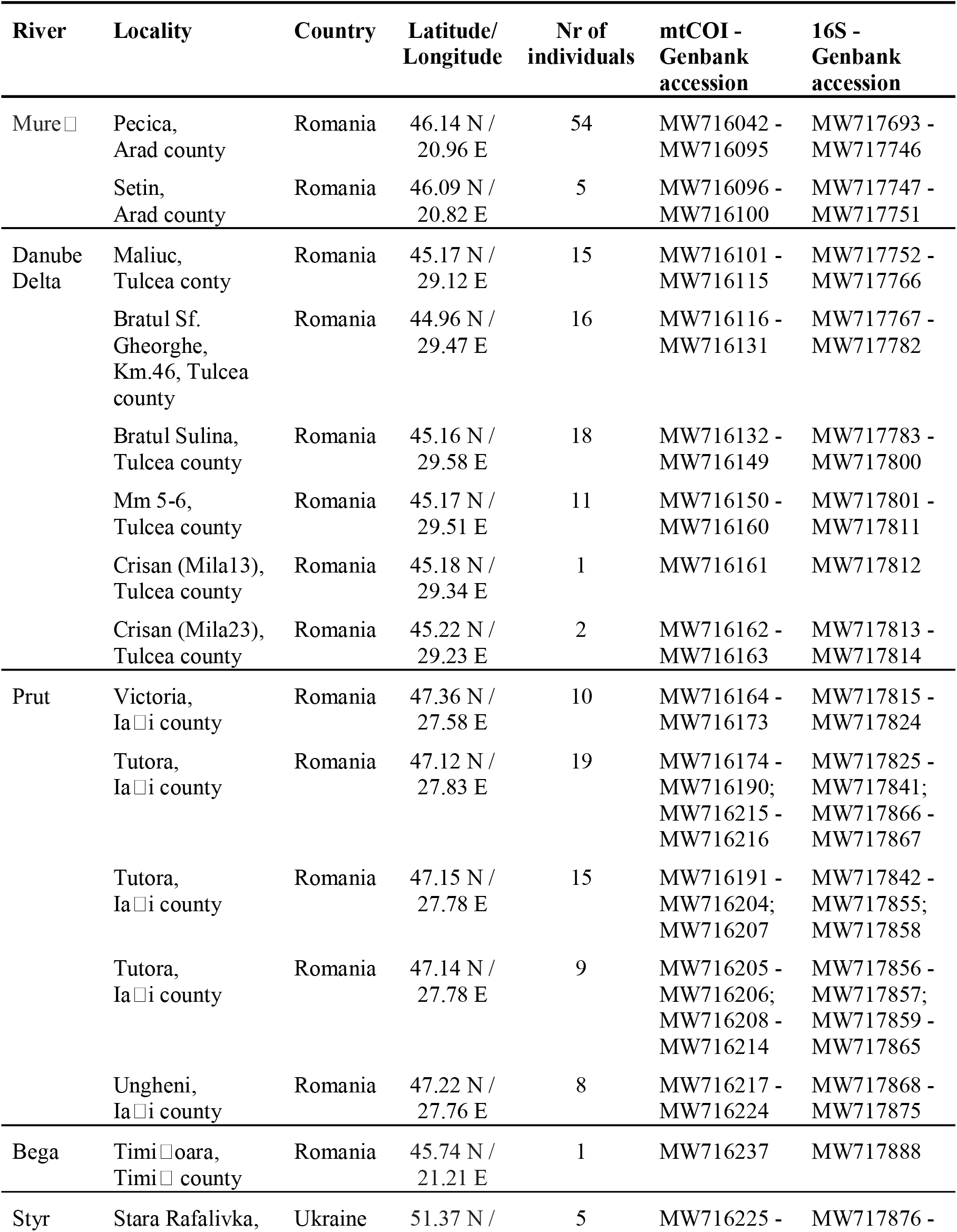
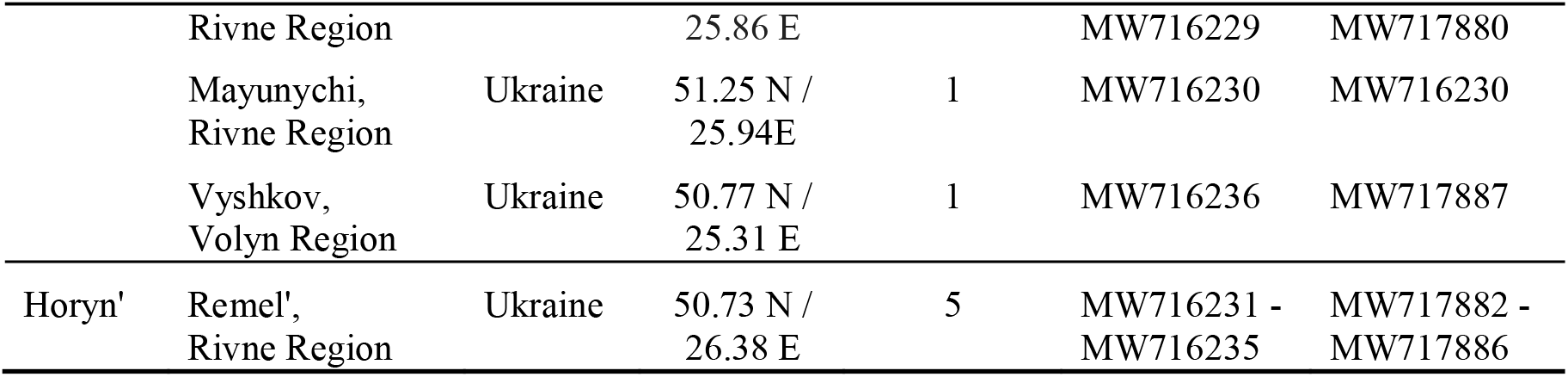
Collection site locations, number of studied specimens, and the GenBank accession codes of the *mt*COI and 16S sequences.

**Figure 2.**
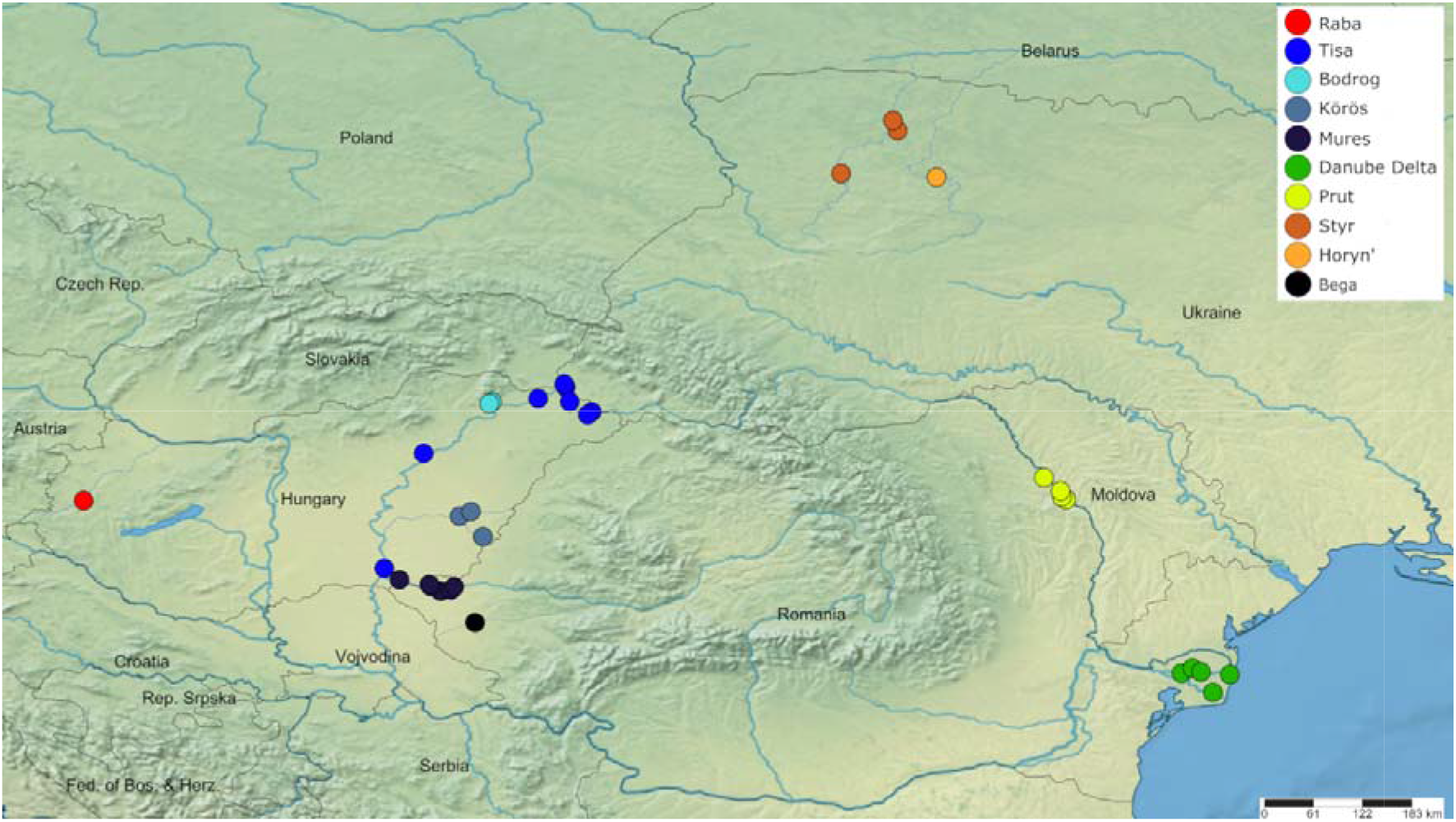
Map showing the collection sites of the individuals used in this study. Only the individuals from Romania (Mure□, Danube Delta, Prut and Bega) and Ukraine (Styr and Horyn’) were collected for this study. The locations from Hungry (Rába, Tisa, Bodrog, Körös and Mure□) are described by Bálint *et al*. (2012).

Two hundred forty five sequences of both markers, generated from the previous study, corresponding to the Hungarian populations of the studied species were also downloaded from the NCBI database (accession numbers, *mt*COI: HE650151 – HE650395; 16S: HE650420 – HE650664). The combined dataset of the current study and from the available Hungarian sequences was used for all further analysis.

### Estimates of genetic diversity

The number of haplotypes and of polymorphic sites (S), the haplotype (Hd) and nucleotide diversity (π) of the *mt*COI, the 16S and the concatenated data sets were calculated in DnaSp 6 (Rozas *et al*., 2017).

### Population structure and patterns of diversity

The *mt*COI and 16S datasets were checked against conflicting phylogenetic information based on the topology of the Neighbor-Joining trees generated with 10000 bootstrap replicates in Mega X (Kumar *et al*., 2018) – data not shown. A Median-Joining (MJ) haplotype network was generated for the concatenated dataset using PopArt 1.7 (Leigh & Bryant, 2015). The haplotypes in the network were grouped based on the rivers they were collected on. A spatial clustering of individuals was also implemented in BAPS 6 (Bayesian Analysis of Population Structure) (Corander *et al*., 2008). This model combines sample locations with likelihood of the genetic data (Cheng *et al*., 2013). The analysis was performed using several runs with default parameters to identify the correct number of partitions.

The genetic differentiation between rivers was estimated with an exact test of population differentiation (ETPD) based on haplotype frequencies (Raymond & Rousset, 1995), and with the pairwise *F*_*ST*_ values using Arlequin 3.5 (Excoffier & Lischer, 2010). ETPD was ran with 100000 Markov Chain steps and 10000 dememorization steps, and *F*_*ST*_ values based on pairwise distances were calculated with 10000 permutations.

The genetic population structure was examined with the hierarchical analysis of the molecular variance (AMOVA) with Arlequin 3.5 (Excoffier & Lischer, 2010), using pairwise distances and 10000 permutations. The hierarchical grouping consisted of the individuals, at the lowest level, grouped together based on the collection sites, and assigned to the corresponding rivers, representing the highest hierarchical level.

### Demographic history and gene flow

The recent demographic history was explored with two approaches. First, Tajima’s *D* index (Tajima, 1989) and Fu’s *F*s test (Fu, 1997) were calculated using Arlequin 3.5 (Excoffier and Lischer, 2010), with 10,000 simulated samples. As a second test, a mismatch distribution analysis was employed in Arlequin 3.5 (Excoffier and Lischer, 2010) under a model of sudden expansion, with 10000 bootstrap replicates. This analysis calculates the frequency distribution of pairwise differences between haplotypes in a population and compares it to the simulated models fitted to the data. A unimodal distribution shows that a lineage has undergone recent population expansion, while a multimodal suggests a constant population size or geographical subdivision (Marjoram & Donnelly, 1994). The appropriateness of this model was evaluated by the sum of squared deviations (SSD) and Harpending’s raggedness index (RI) (Harpending, 1994). Both approaches were used for the whole dataset and for the separate rivers.

## Results

### Estimates of genetic diversity

The 196 sequences generated by this study showed *S* = 30 variable sites for the *mt*COI, resulting in 32 haplotypes with a haplotype diversity of *Hd =* 0.812 and a nucleotide diversity of π *=* 0.0037. The 16S region had *S* = 39 variable sites that led to 42 haplotypes with a haplotype diversity of *Hd =* 0.780 and a nucleotide diversity of π *=* 0.0028. The concatenated sequences showed *S* = 69 variable sites, *Hd =* 0.900 haplotype diversity and of π *=* 0.0032 nucleotide diversity. The combined datasets included a total of 441 sequences for each marker. The *mt*COI alignment showed a high genetic diversity, with *S* = 48 variable sites, resulting in 57 haplotypes, of which three had high frequencies (represented by 81, 133 and 133 individuals) and 41 were unique (represented by only one specimen). The haplotype diversity was *Hd =* 0.7846 and the nucleotide diversity was π *=* 0.0035. In the case of the 16S alignment, the number of variable sites was *S =* 64, resulting in 86 haplotypes, showing a similar pattern to that of the *mt*COI data, with three frequent (represented by 88, 115 and 138 sequences) and 71 unique haplotypes (represented by only one specimen). The haplotype diversity was *Hd =* 0.7955 and the nucleotide diversity was π *=* 0.0031. The concatenated dataset showed similar high genetic diversity and low divergence. The number of polymorphic sites was *S =* 112 and the number of haplotypes was 148, also with three frequent haplotypes, designated as H1, H2 and H3 (represented by 60, 88 and 100 individuals respectively), and 123 unique haplotypes (represented by only one specimen) (Figure 3). The other haplotypes are represented by 2 to 6 specimens. The haplotype diversity was *Hd =* 0.8908 and the nucleotide diversity was π *=* 0.0033. Only one individual was collected from the Bega River and corresponded to H3. This river was not used in further analysis due to the lack of information. The two rivers from Ukraine, Styr (7 individuals) and Horyn’ (5 individuals), were considered as one group as tributaries of Pripyat River.

**Figure 3.**
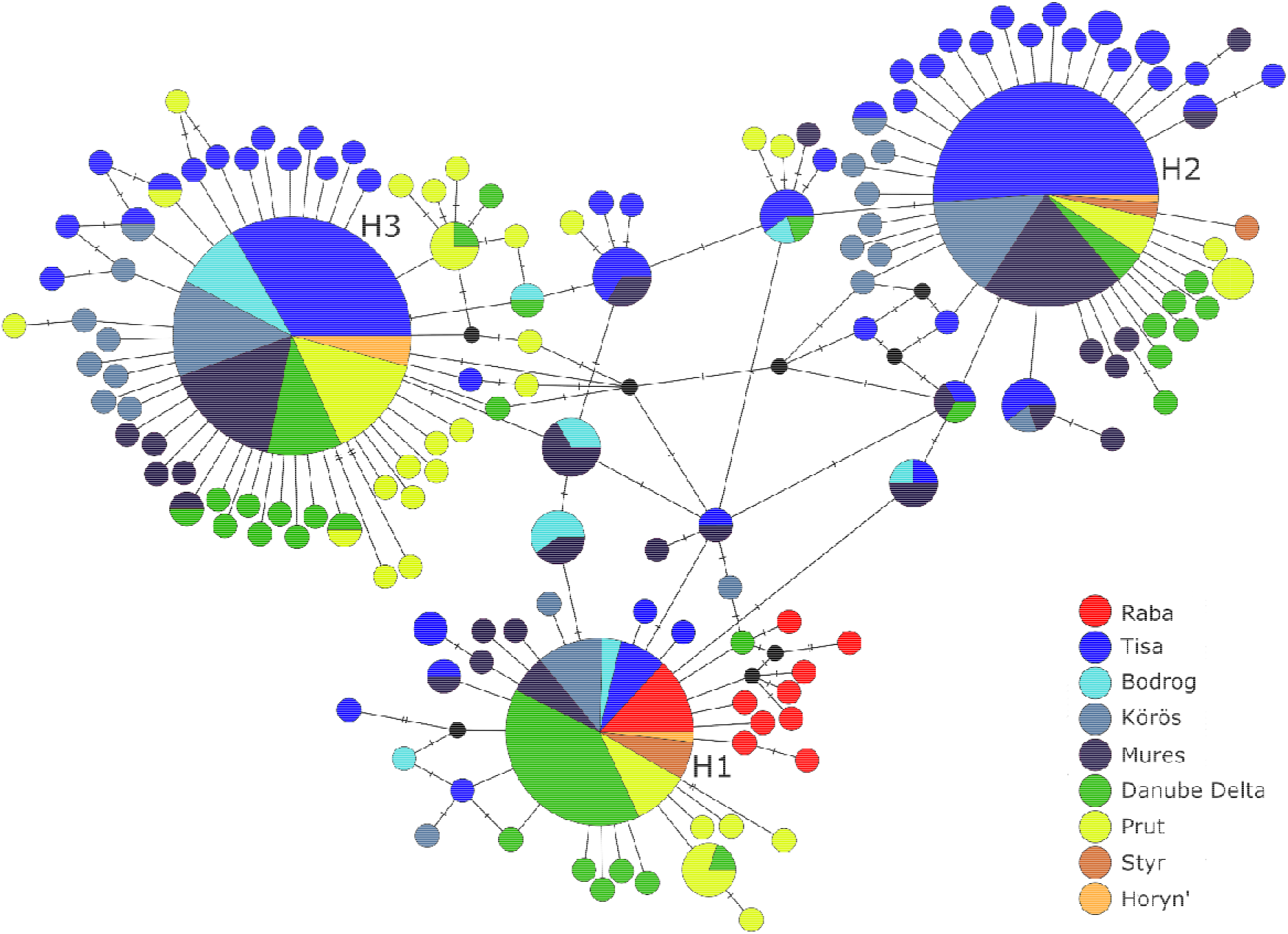
Median-Joining haplotype network generated for the concatenated dataset using PopArt 1.7. Each circle represents a unique haplotype and circle size is proportional to the number of samples observed for that haplotype. The number of mutations is represented by hatch marks on the lines. Colors correspond to different rivers. H1, H2 and H3 correspond to the three major haplotypes discussed in the text.

### Population structure and patterns of diversity

The MJ network shows no geographic differentiation, the three frequent haplotypes were represented by individuals from almost every river. Two of them (H2 and H3) are more frequent in the Tisa River and its tributaries (Bodrog, Körös rivers and Mure□), but are represented also by individuals from the Danube Delta, Prut River and the tributaries of Pripyat (Figure 3). The third major haplotype (H1) is present more frequently in specimens from the Danube Delta, followed by the Tisa catchments, and individuals from Rába and from Prut rivers. Besides the three frequent haplotypes, the Tisa and its tributaries share four haplotypes with the Danube Delta and one with Prut. The Delta and Prut have three additional common haplotypes (Figure 3).

The Bayesian Analysis of Population Structure defined two groups (optimal partition, log(likelihood) = -3489.23), also without showing any clear geographic pattern. The first group includes H1, H3 and their satellite haplotypes corresponding to each river. The second group has H2 in the central position, and it does not contain specimens collected from the Rába River. Although both groups are dominated by specimens from the Tisa and its tributaries, the majority of individuals collected in the Danube Delta and from the Prut River belong to second group (82.53% of the Delta and 81.96% of the Prut specimens).

The two lineages detected by BAPS were also confirmed by the significant differentiation showed by the pairwise *F*_*ST*_ = 0.556; *p* < 0.001 and ETPD *p* < 0.001. Population pairwise *F*_*ST*_ calculations showed that the Rába River is significantly differentiated from all other rivers, with an average of *F*_*ST*_ = 0.310 (Table 2). The lowest pairwise difference was shown between the Rába River and the Danube Delta (*F*_*ST*_ = 0.155, *p* < 0.001). The ETPD shows no significant differentiation between these two rivers (*p =* 0.19; Table 2). The *F*_*ST*_ values were significant when the Tisa River was compared with the Danube Delta (*F*_*ST*_ = 0.128; *p* < 0.001) and Prut (*F*_*ST*_ = 0.111; *p* < 0.001). These differences were supported also by the significant differentiation values (*p* < 0.001) of the ETPD (Table 2). Interestingly, there was significant pairwise difference and differentiation between the Tisa and Bodrog rivers (*F*_*ST*_ = 0.174, *p* < 0.001; ETPD: *p* < 0.05). A lower, but significant difference was observed between the Delta and Prut (*F*_*ST*_ = 0.064, *p* < 0.001), with strong support for the differentiation at *p* < 0.001 (Table 2).

**Table 2.** Genetic differentiation of populations from the different rivers. Pairwise *Fst* values (lower left) and significant ETPD (upper right) results of extant populations. Bold values are significant at: ^**^<0.001 and ^**^<0.01.

|  | (1) | (2) | (3) | (4) | (5) | (6) | (7) | (8) |
| --- | --- | --- | --- | --- | --- | --- | --- | --- |
| (1) Rába |  | + | + | + | + | - | + | - |
| (2) Tisa | **0.372** ** |  | - | - | - | + | + | - |
| (3) Bodrog | **0.382** ** | **0.174** ** |  | - | - | - | - | - |
| (4) Körös | **0.333** ** | 0.002 | **0.107** * |  | - | + | - | - |
| (5) Mureș | **0.346** ** | 0.001 | **0.117** * | -0.009 |  | + | + | - |
| (6) Danube Delta | **0.155** ** | **0.128** ** | 0.060 | **0.073** * | **0.082** ** |  | + | - |
| (7) Prut | **0.312** ** | **0.111** ** | 0.022 | **0.054** * | **0.069** ** | **0.064** ** |  | - |
| (8) Pripyat basin | **0.276** ** | 0.006 | 0.089 | -0.028 | -0.022 | -0.011 | 0.026 |  |

The analysis of molecular variance showed that most of the variance was found within collection sites (89.20%, *F*_*ST*_ = 0.108, *p* < 0.001), followed by the variance between rivers (7.85%, *F*_*CT*_ = 0.078, *p* < 0.001). The lowest variation was found among collection sites within the different rivers (2.95%, *F*_*SC*_ = 0.032, *p <* 0.05).

### Demographic history and gene flow

The analysis of the demographic history for the whole dataset shows significant departure from the equilibrium. Both Tajima’s *D* index and Fu’s *F*s test showed negative values with significant support (Tajima’s *D* = -2.393, *p* < 0.001; Fu’s *F*s = -25.762, *p* < 0.001), and the mismatch distribution plot (Figure S1) fits well with the sudden population expansion model (SSD = 0.0119, *p* = 0.062; Raggedness index = 0.028, *p* = 0.16). The two BAPS lineages also showed significant departure from equilibrium, based on the neutrality tests, and recent expansion, based on the mismatch distribution results (Table 3; Figure S1). Similar results were observed on a regional scale, where Tajima’s *D* and Fu’s *F*s showed negative values and significant departures from equilibrium for all rivers, except for the Bodrog River and the Pripyat tributaries (Table 3). The mismatch distribution plots, SSD and Raggedness index values support the sudden expansion model for all rivers, except the Pripyat tributaries (Table 3; Figure S1).

**Table 3.** Results of mismatch distribution and neutrality tests for the whole dataset, the two linages identified by BAPS, and for populations from different rivers.

|  | Mismatch distribution |  |  |  | Test of selective neutrality |  |  |  |
| --- | --- | --- | --- | --- | --- | --- | --- | --- |
|  | SSD | *p* | RI | *p* | Tajima's D | *p* | Fu's Fs | *p* |
| **All** | 0.011 | 0.062 | 0.028 | 0.162 | -2.393 | 0.000 | -25.762 | 0.000 |
| **BAPS1** | 0.005 | 0.573 | 0.0210 | 0.752 | -2.433 | 0.000 | -26.570 | 0.000 |
| **BAPS2** | 0.001 | 0.230 | 0.056 | 0.106 | -2.676 | 0.000 | -29.892 | 0.000 |
| **Rába** | 0.049 | 0.078 | 0.199 | 0.046 | -1.818 | 0.021 | -5.399 | 0.0008 |
| **Tisa** | 0.015 | 0.290 | 0.032 | 0.461 | -1.801 | 0.008 | -26.433 | 0.000 |
| **Bodrog** | 0.006 | 0.736 | 0.029 | 0.908 | -0.258 | 0.442 | -1.556 | 0.152 |
| **Körös** | 0.021 | 0.093 | 0.047 | 0.191 | -1.515 | 0.044 | -15.278 | 0.000 |
| **Mureș** | 0.015 | 0.115 | 0.041 | 0.183 | -1.335 | 0.068 | -23.833 | 0.000 |
| **Delta** | 0.016 | 0.317 | 0.034 | 0.452 | -1.938 | 0.008 | -23.071 | 0.000 |
| **Prut** | 0.008 | 0.054 | 0.026 | 0.199 | -1.829 | 0.012 | -25.914 | 0.000 |
| **Pripyat** | 0.120 | 0.009 | 0.284 | 0.043 | 1.159 | 0.880 | 1.578 | 0.809 |

## Discussion

### The phylogeographic pattern of P. longicauda

The results of this study show a pattern similar to that observed by Bálint *et al*. (2012), with high haplotype diversity and low divergence of individuals collected from Romania and Ukraine, as well as of the combined datasets. Both markers used in this study have a high mutation rate and were successfully used in intraspecific level phylogeographic analysis (Takenaka *et al*., 2019; Hrivniak *et al*., 2020). However, for *P. longicauda* populations both markers lack sufficient phylogenetic resolution. The observed high diversity and low divergence may be a result of incomplete linage sorting, interbreeding of individuals from different lineages or migration waves (Baggiano *et al*., 2011; Sworobowicz *et al*., 2020).

The biogeography of the Western Palearctic was proven to be more complex than the classical theory of the three Mediterranean glacial refugia, as the genetic analysis of several different groups supported the presence of many extra-Mediterranean refugia throughout Europe (Schmitt & Varga, 2012; Wattier *et al*., 2020). Studies focusing on the Danube catchment identified the Ponto-Caspian Region and the Pannonian Basin as extra-Mediterranean refugia areas for many vertebrate and invertebrate taxa (Bănăduc *et al*., 2016; Vörös *et al*., 2016; Csapó *et al*., 2020). There are several established theories/paradigms regarding the genetic diversity of a region. On one hand, a refugia is considered to be the most diverse region, a “hot spot”, with the expansion of the range leading to the loss of diversity toward the edge, as a result of successive founder events (Avise *et al*., 1987; Hewitt, 2004). On the other hand, a region could show a high genetic diversity, due to secondary accumulation of dispersing lineages of different geographic origin and evolutionary history, a “melting pot” (Petit *et al*., 2003).

The Pontic region is considered an important diversification center and refugia area for many species, and the post-glacial upstream recolonization of North-western Europe through the Danube Basin is a well-established paradigm of the freshwater zoogeography (Bănărescu, 1991; Varga, 2010). This pattern would explain the presence of the same *P. longicauda* haplotype (H1) in the Tisa and its tributaries, as well as in the Rába River and, based on the short (196 bp) sequences, the Rhine River. The colonization of the Prut River, as the closest Danube tributary, and the two Ukrainian rivers (Styr and Horyn’) from the Pontic region is also plausible, as the Pripyat River is a tributary of the Dnieper (or Dnipro) River, which is, together with Southern Bug (or Pivdennyi Buh) and Dniester (or Dnister) rivers, a potential migration corridor for aquatic biota from the Black sea coast to North-eastern Europe (Bij de Vaate *et al*., 2002; Sworobowicz *et al*., 2020).

The Pannonian Basin and the middle Danube catchment is considered an important cumulative refugia for several different faunal types, due to the development of a boreal forest-steppe in the region, during the Pleistocene (Varga, 2010). This region is considered to have harbored *P. longicauda* in at least one refugium (Bálint *et al*., 2012). The authors argue that the explanation for the strongly diverged private haplotypes in the Rába River is, that there were two close or overlapping refugia in the middle Danube catchment. The haplotype network generated based on the combined datasets shows one unique haplotype from the Rába River linked to another haplotype from the Danube Delta, which is linked to H1, the most frequent haplotype from the Danube Delta. The close connection of the Rába River with the Danube Delta is also confirmed by the low pairwise difference value and the result of the ETPD, which showed no significant differentiation.

The MJ network reflects the low nucleotide diversity, showing no clear phylogeographic pattern. Three frequent haplotypes, present in every studied river, dominate the star-like network, but there are also several unique haplotype for each of these rivers. The Tisa and its tributaries are the most present, being represented by the largest number of shared and unique haplotypes. This is in concordance with the cumulative refugia role and the “melting pot” hypothesis (Dufresnes *et al*., 2016). In the Danube Delta H1 is the most frequent, but all three major haplotypes are present, and a high number of unique satellite haplotypes are linked to each of them. All three frequent haplotypes are present in the Prut River, but the satellite haplotypes are mostly linked to the H3. The Pripyat tributaries show the same pattern, but a more intensive sampling is needed in the region.

The BAPS analysis showed two differentiated lineages, confirmed also by the ETPD and *F*_*ST*_ values. This could reflect a possible Pannonian refugia somewhere in the middle-Danube region, as suggested by Bálint *et al*. (2012), and a Pontic refugia, which would be in concordance with the biogeographic pattern shown by several aquatic biota (Antal *et al*., 2016; Vörös *et al*., 2016; Sworobowicz *et al*., 2020). However, the high haplotype diversity, the low genetic differentiation, the significant differences (*F*_*ST*_) and differentiations (ETPD) between the rivers, confirmed also by the relatively high variance between them (AMOVA), leads us to speculate the possibility of an interconnected microrefugia network between the Pannon and the Pontic region, making the species a Ponto-Pannonian element. During the interglacial periods, and after the LGM, the melting of the ice sheets led to more complex hydrological networks (Panin *et al*., 2020), which would have allowed the dispersal of individuals from different microrefugia, leading to the interbreeding of the different lineages. The concept of a vast network of microrefugia, with favorable habitat for small invertebrates was also suggested by Sworobowicz *et al*. (2020), and the large number of studies that found evidence of refugia for different taxonomic groups throughout Europe, could also be considered as evidence for such a network.

### Conservation implications

River regulations, damming, hydropower plant construction and pollution has drastically altered aquatic habitats of European rivers in the last centuries, leading to a long term process of decline in freshwater insect biodiversity (Assandri, 2021; Jähnig *et al*., 2021), especially sensitive groups like Ephemeroptera, Plecoptera and Trichoptera (Graf *et al*.,2015; Krno *et al*., 2018). However, in the recent years water quality of the Danube showed improvement (Liska, 2015; Stoica *et al*., 2019) as a result of implementation of the EU Water Framework Directive and of the EU Floods Directive by the International Commission for the Protection of the Danube River (ICPDR), facilitating the return or recovery of some sensitive species. *Ephoron virgo* (Olivier, 1791), an European burrowing filter-feeding mayfly, became a symbol for the recovery of polluted rivers after being reported from the Danube basin by several studies (Vidinova & Russev, 1997; Kovács *et al*., 2001; Adámek *et al*., 2007; Marković *et al*., 2017).

The decline of *P. longicauda*, a habitat specialist of clayey substrates, was the result of organic pollution and riverbank regulations that started at the end of the 19^th^ century (Russev, 1987; Tittizer *et al*., 2008). However, Bálint *et al*. (2012) suggested the possibility of cryptic survival of this species on its formerly vast distribution range. The phylogeographic results of the present study indicate that the presence of the analyzed populations is not the result of recent, secondary recolonization from the Pannon region, confirming the survival and expansion of cryptic local lineages. The physical and hydrological characteristics of the lower section of the Danube River (Romania and Bulgaria) and the Danube Delta show good conditions in large sectors, being only slightly or moderately regulated, with nearly natural banks or small sections of reinforced banks, and with floodplains of high or moderate ecological value (Schwarz, 2015). These conditions could have also facilitated the survival of the species and its recent expansion in the region. The natural habitats of the Prut River are also well preserved (Vartolomei, 2012), allowing them to be designated as a Natura 2000 (SCI and SPA) sites and IUCN protected areas (Category IV: Habitat/Species Management Area – Nature Reserve, and Category V: Protected Landscape/Seascape – Natural Park), making the presence of *P. longicauda* populations possible.

The real natural phenomena of the mass swarming of the *P. longicauda* adults got important social interest in the past, because this species was widely used as bait for fishing. It was popularly known under various names: “oeveraas” and “haft” in the Netherlands, “Spork-Oese”, “Sprock”, “Spaargoos”, “Spaargaanse” in Germany, “tiszavirág” in Hungary, “gandatsi” for larvae and “rusalki”, or “karchani” for adults in Bulgaria (Russev, 1987). In Romania it is popularly known as the “flower of the rivers” or “rusalii” in the Danube Delta and Mure□, and on the Prut River, under the name of “vetrică”. In the present digitalized information network, including various social media platforms, there is a lot of information about this species, either as a tourist attraction or as important information among fishermen, because it is still used as bait for fishing in Romania.

Phylogeographic data can contribute to clarify the conservation status and long term population management of some endemic or endangered species. A good example is the Hungarian Lilac (*Syringa josikae* J. Jacq. ex Rchb. f.), an endangered plant species, endemic to the Apuseni Mts. (Romania) and Eastern Carpathians (Ukraine). This species had only sporadic data from its range, and lacked any focused management of its highly reduced populations, but the situation was change by the contribution of Lendvay *et al*. (2016), from data deficient (DD) to endangered (EN) (Höhn and Lendvay, 2018).

A similar situation can be observed for *Palingenia longicauda*, as its current status in Romania is not evaluated (DD – Data Deficient), and even if in some EU states it is considered highly endangered (Hungary, Ukraine), presently it is not included in the IUCN Red list of endangered species. Based on our results, we will propose to change the IUCN status of this species from insufficient date (DD) to endangered (EN), and its inclusion in the Romanian National Red Lists. This will help the development of an effective management for the sustainable conservation of this species, and will have an important impact on the conservation of *P. longicauda* in the larger European context.

## Conclusions

*Palingenia longicauda*, considered extinct from the major part of its distribution (except the Tisa catchment), was reported in the last two decades as present in the Danube river in Hungary (Málnás *et al*., 2016), in the Danube Delta in Romania and Ukraine (Soldán *et al*., 2009; Bulánková *et al*., 2013; Pavel *et al*., 2019; Afanasyev *et al*., 2020), and in the Prut River in the Republic of Moldova (Munjiu, 2018). An important result of this study is that it confirms the presence of viable populations in the Danube Delta and on the Prut River in Romania, and that further shows the species presence on the Mure□ and Bega rivers from Romania, and on the Styr and Horyn’ rivers in Ukraine. This first genetic study of the newly identified populations suggest the need of a more intensive survey, to help identify and protect these cryptic populations on the species whole former distribution.

The recent recovery of the species may be related to the improvement of water quality as a result of the international effort in conservation of freshwater ecosystems, supporting the necessity of the implementation EU Water policies. However, continuous transformations caused by river bed diggings and regulations, or environmental catastrophe threats (like the repeated cyanide pollution of the Tisa River, from 2000 onward) show the need of a stronger cooperation between science and society to maintain the unique biodiversity of large rivers in the Danube River catchment area.

## Acknowledgements

This work was supported by two grant of the Romanian Ministry of Education and Research, CNCS - UEFISCDI, project numbers PN-III-P1-1.1-PD-2019-0829; nr. PD91/2020 and PN-III-P2-2.1-PED-2019-0214; nr. 476PED/2020. The work of Martynov A.V. was partially supported by SIU (Norwegian Centre for International Cooperation in Education) grant CPEA-LT-2016/10140 to Vladimir Gusarov (Natural History Museum, University of Oslo). During the study and preparation of the manuscript Szabó E. received financial support through the Collegium Talentum Program of Hungary. The project also got support from the European Cooperation in Science and Technology (COST) Action DNAqua-Net (CA15219) Working Group 1, led by Torbjørn Ekrem and Fedor Čiampor.

---

Figure S1. Mismatch distribution histograms, for the whole dataset (All), the two groups identified by the BAPS (BAPS1 and BABS2), and for populations from each river. Bars indicate the observed values and black lines show the expected distribution under the sudden expansion model.

## Notes

### Competing Interest Statement

The authors have declared no competing interest.

## References

Adámek, Z., Andreji, J. & Gallardo, J. M. (2007) Food habits of four bottom-dwelling gobiid species at the confluence of the Danube and Hron Rivers (South Slovakia). International Review of Hydrobiology, 92, 554–563.

Afanasyev, S., Lyashenko, A., Iarochevitch, A., Lietytska, O., Zorina-Sakharova, K. & Marushevska, O. (2020) Pressures and Impacts on Ecological Status of Surface Water Bodies in Ukrainian Part of the Danube River Basin. Human Impact on Danube Watershed Biodiversity in the XXI Century (ed. by Bănăduc, D., Curtean-Bănăduc, A., Pedrotti, F., Cianfaglione, K. & Akeroyd, J.R.), pp. 327–358. Springer, Cham.

Akimov, I.A. (2009) Red Data Book of Ukraine. Animals. 3rd ed. Global Consulting, Kyiv, Ukraine.

Albert, J.S., Destouni, G., Duke-Sylvester, S.M., Magurran, A.E., Oberdorff, T., Reis, R.E., Winemiller, K.O. & Ripple, W. J. (2021) Scientists’ warning to humanity on the freshwater biodiversity crisis. Ambio, 50, 85–94.

Alexander, L.C., Hawthorne, D.J., Palmer, M.A. & Lamp, W.O. (2011) Loss of genetic diversity in the North American mayfly Ephemerella invaria associated with deforestation of headwater streams. Freshwater Biology, 56, 1456–1467.

Andrikovics, S., Fink, T.J. & Cser, B. (1992) Tisza mayfly monograph. Booklet of Tisza Club, 2, 9–35.

Antal, L., László, B., Kotlík, P., Mozsár, A., Czeglédi, I., Oldal, M., Kemenesi, G., Jakab, F. & Nagy, S.A. (2016) Phylogenetic evidence for a new species of Barbus in the Danube River basin. Molecular Phylogenetics and Evolution, 96, 187–194.

Assandri, G. (2021) Anthropogenic-driven transformations of dragonfly (Insecta: Odonata) communities of low elevation mountain wetlands during the last century. Insect Conservation and Diversity. 14, 26–39.

Avise, J.C., Arnold, J., Ball, R.M., Bermingham, E., Lamb, T., Neigel, J.E., Reeb, C.A. & Saunders, N.C. (1987) Intraspecific Phylogeography: The Mitochondrial DNA Bridge Between Population Genetics and Systematics. Annual Review of Ecology and Systematics, 18, 489–522.

Babic, W., Branicki, W., Crnobrnja-Isailović, J., Cogălniceanu, D., Sas, I., Olgun, K., Poyarkov, N.A., Garcia-París, M. & Arntzen, J.W. (2005) Phylogeography of two European newt species — discordance between mtDNA and morphology. Molecular Ecology, 14, 2475–2491.

Baggiano, O., Schmidt, D.J., Sheldon, F. & Hughes, J.M. (2011) The role of altitude and associated habitat stability in determining patterns of population genetic structure in two species of Atalophlebia (Ephemeroptera: Leptophlebiidae). Freshwater Biology, 56, 230–249.

Bálint, M., Málnás, K., Nowak, C., Geismar, J., Váncsa, É., Polyák, L., Lengyel, S. & Haase, P. (2012) Species history masks the effects of human-induced range loss - unexpected genetic diversity in the endangered giant mayfly Palingenia longicauda. PLoS ONE, 7, e31872–e31880.

Bănăduc, D., Rey, S., Trichkova, T., Lenhardt, M. & Curtean-Bănăduc, A. (2016) The Lower Danube River-Danube Delta-North West Black Sea: A pivotal area of major interest for the past, present and future of its fish fauna - A short review. Science of the Total Environment, 545–546, 137–151.

Bănărescu, P. (1991) Zoogeography of Fresh Waters: Distribution and Dispersal of Freshwater Animals in North America and Eurasia. AULA-Verlag, Wiesbaden, Germany.

Bauernfeind, E. & Soldán, T. (2012) The Mayflies of Europe (Ephemeroptera). Apollo Books, Ollerup, Denmark.

Bij de Vaate, A., Jazdzewski, K., Ketelaars, H.A.M., Gollasch, S. & Van der Velde, G. (2002) Geographical patterns in range extension of Ponto-Caspian macroinvertebrate species in Europe. Canadian Journal of Fisheries and Aquatic Sciences, 59, 1159– 1174.

Bulánková, E., Beracko, P. & Derka, T. (2013) Occurrence of protected species (Gomphus flavipes, Odonata and Palingenia longicauda, Ephemeroptera) in the Danube River and its deltas (Romania, Slovakia)’, Scientific Annals of the Danube Delta Institute, 19, 21–24.

Carrizo, S.F., Lengyel, S., Kapusi, F., Szabolcs, M., Kasperidus, H.D., Scholz, M., Markovic, D., Freyhof, J., Cid, N., Cardoso, A.C. & Darwall, W. (2017) Critical catchments for freshwater biodiversity conservation in Europe: identification, prioritisation and gap analysis. Journal of Applied Ecology, 54, 1209–1218.

Cheng, L., Connor, T.R., Sirén, J., Aanensen, D.M. & Corander, J. (2013) Hierarchical and spatially explicit clustering of DNA sequences with BAPS software. Molecular Biology and Evolution, 30, 1224–1228.

Coleman, R.A., Gauffre, B., Pavlova, A., Beheregaray, L.B., Kearns, J., Lyon, J., Sasaki, M., Leblois, R., Sgro, C. & Sunnucks, P. (2018) Artificial barriers prevent genetic recovery of small isolated populations of a low-mobility freshwater fish. Heredity, 120, 515–532.

Collen, B., Whitton, F., Dyer, E.E., Baillie, J.E.M., Cumberlidge, N., Darwall, W.R.T., Pollock, C., Richman, N.I., Soulsby, A.-M. & Böhm, M. (2014) Global patterns of freshwater species diversity, threat and endemism. Global Ecology and Biogeography, 23, 40–51.

Corander, J., Sirén, J. & Arjas, E. (2008) Bayesian spatial modeling of genetic population structure, Computational Statistics, 23, 111–129.

Csapó, H., Krzywoźniak, P., Grabowski, M., Wattier, R., Bącela-Spychalska, K., Mamos, T., Jelić, M. & Rewicz, T. (2020) Successful post-glacial colonization of Europe by single lineage of freshwater amphipod from its Pannonian Plio-Pleistocene diversification hotspot. Scientific Reports, 10, 18695.

Darwall, W., Holland, R., Smith, K., Allen, D. & Brooks, E. (2011) Investment for freshwater species. Conservation Letters, 4, 474–482.

Dijkstra, K.D.B., Monaghan, M.T. & Pauls, S.U. (2014) Freshwater Biodiversity and Aquatic Insect Diversification. Annual Review of Entomology, 59, 143–163.

Dufresnes, C., Litvinchuk, S.N., Leuenberger, J., Ghali, K., Zinenko, O., Stöck, M. & Perrin, N. (2016) Evolutionary melting pots: a biodiversity hotspot shaped by ring diversifications around the Black Sea in the Eastern tree frog (Hyla orientalis). Molecular Ecology, 25 4285–4300.

Dupuis, J.R., Geib, S.M., Schmidt, C. & Rubinoff, D. (2020) Genomic-wide sequencing reveals remarkable connection between widely disjunct populations of the internationally threatened bog buck moth. Insect Conservation and Diversity, 13, 495–500.

Excoffier, L. & Lischer, H.E.L. (2010) Arlequin suite ver 3.5: a new series of programs to perform population genetics analyses under Linux and Windows. Molecular Ecology Resources, 10, 564–567.

Fu, Y.X. (1997) Statistical tests of neutrality of mutations against population growth, hitchhiking and background selection. Genetics, 147, 915–925.

Graf, W., Leitner, P. & Pletterbauer, F. (2015) Short overview on the benthic macroinvertebrate fauna of the Danube River. The Handbook of Environmental Chemistry, 39, The Danube River Basin. (ed. by Liška, I.), pp. 287–315. Heidelberg: Springer-Verlag, Berlin, Germany.

Hall, T.A. (1999) BioEdit: a user-frendly biological sequence alignment editor and analysis program foe Windows 95/98/NT. Nucleic Acid Symposium Series, 41, 95–98.

Harpending, H.C. (1994) Signature of ancient population growth in a low-resolution mitochondrial DNA mismatch distribution. Human Biology, 66, pp. 591–600.

Haybach, A. (1998) Die Eintagsfliegen (Insecta: Ephemeroptera) von Rheinland-Pfalz Zoogeographie, Faunistik, Ökologie, Taxonomie und Nomenklatur - Unter besonderer Berücksichtigung der Familie Heptageniidae und unter Einbeziehung der übrigen aus Deutschland bekannten Arten. Dissertation, Mainz University.

Hein, T., Schwarz, U., Habersack, H., Nichersu, I., Preiner, S., Willby, N. & Weigelhofer, G. (2016) Current status and restoration options for floodplains along the Danube River. Science of the Total Environment, 543, 778–790

Hewitt, G. (1999) Post-glacial re-colonization of European biota. Biological Journal of the Linnean Society, 68, 87–112.

Hewitt, G. (2004) Genetic consequences of climatic oscillations in the Quaternary. Phyolsophical Travsactions of The Rolyal Society, 359, 183–195.

Höhn, M. & Lendvay, B. (2018) Syringa josikaea. The IUCN Red List of Threatened Species 2018, e.T162267A99428926.

Hrivniak, Ľ., Sroka, P., Bojková, J., Godunko, R.J., Soldán, T. & Staniczek, A.H. (2020) The impact of Miocene orogeny for the diversification of Caucasian Epeorus (Caucasiron) mayflies (Ephemeroptera: Heptageniidae). Molecular Phylogenetics and Evolution, 146, 106735.

Jähnig, S.C., Baranov, V., Altermatt, F., Cranston, P., Friedrichs-Manthey, M., Geist, J., He, F., Heino, J., Hering, D., Hölker, F., Jourdan, J., Kalinkat, G., Kiesel, J., Leese, F., Maasri, A., Monaghan, M.T., Schäfer, R.B., Tockner, K., Tonkin, J.D. & Domisch, S. (2021) Revisiting global trends in freshwater insect biodiversity. Wiley Interdisciplinary Reviews: Water, 8, e1506–e1510.

Johnson, M., Zaretskaya, I., Raytselis, Y., Merezhuk, Y., McGinnis, S. & Madden, T.L. (2008) NCBI BLAST: a better web interface. Nucleic Acids Research, 3, pp. 5–9.

Jourdan, J., Plath, M., Tonkin, J.D., Ceylan, M., Dumeier, A.C., Gellert, G., Graf, W., Hawkins, C.P., Kiel, E., Lorenz, A.W., Matthaei, C.D., Verdonschot, P.F.M., Verdonschot, R.C.M. & Haase, P. (2018) Reintroduction of freshwater macroinvertebrates: challenges and opportunities. Biological Reviews, 94, 368–387.

Kovács, T., Juhasz, P. & Turcsányi, I. (2001) Ephemeroptera, Odonata and Plecoptera larvae from the river Tisza (1997-1999). Folia Historico Naturalia Musei Matraensis, 25, 135–143.

Kriska, G., Bernáth, B. & Horváth, G. (2007) Positive polarotaxis in a mayfly that never leaves the water surface: Polarotactic water detection in Palingenia longicauda (Ephemeroptera). Naturwissenschaften, 94, 148–154.

Krno, I., Beracko, P., Navara, T., Šporka, F., Mišíková Elexová, E. (2018) Changes in species composition of water insects during 25-year monitoring of the Danube floodplains affected by the Gabčíkovo waterworks. Environmental Monitoring and Assessment, 190, 412.

Kumar, S., Stecher, G., Li, M., Knyaz, C. & Tamura, K. (2018) MEGA X: Molecular evolutionary genetics analysis across computing platforms. Molecular Biology and Evolution, 35, 1547–1549.

Kuntke, F., de Jonge, N., Hesselsøe, M. & Lund, N. (2020) Stream water quality assessment by metabarcoding of invertebrates. Ecological Indicators. 111, 105982.

Leese, F., Altermatt, F., Bouchez, A., Ekrem, T., Hering, D., Meissner, K., Mergen, P., Pawlowski, J., Piggott, J., Rimet, F., Steinke, D., Taberlet, P., Weigand, A., Abarenkov, K., Beja, P., Bervoets, L., Björnsdóttir, S., Boets, P., Boggero, A., Bones, A., Borja, Á., Bruce, K., Bursić, V., Carlsson, J., Čiampor, F., Čiamporová-Zatovičová, Z., Coissac, E., Costa, F., Costache, M., Creer, S., Csabai, Z., Deiner, K., DelValls, Á., Drakare, S., Duarte, S., Eleršek, T., Fazi, S., Fišer, C., Flot, J.-F., Fonseca, V., Fontaneto, D., Grabowski, M., Graf, W., Guðbrandsson, J., Hellström, M., Hershkovitz, Y., Hollingsworth, P., Japoshvili, B., Jones, J., Kahlert, M., Kalamujic Stroil, B., Kasapidis, P., Kelly, M., Kelly-Quinn, M., Keskin, E., Kõljalg, U., Ljubešić, Z., Maček, I., Mächler, E., Mahon, A., Marečková, M., Mejdandzic, M., Mircheva, G., Montagna, M., Moritz, C., Mulk, V., Naumoski, A., Navodaru, I., Padisák, J., Pálsson, S., Panksep, K., Penev, L., Petrusek, A., Pfannkuchen, M., Primmer, C., Rinkevich, B., Rotter, A., Schmidt-Kloiber, A., Segurado, P., Speksnijder, A., Stoev, P., Strand, M., Šulčius, S., Sundberg, P., Traugott, M., Tsigenopoulos, C., Turon, X., Valentini, A., van der Hoorn, B., Várbíró, G., Vasquez Hadjilyra, M., Viguri, J., Vitonytė, I., Vogler, A., Vrålstad, T., Wägele, W., Wenne, R., Winding, A., Woodward, G., Zegura, B. & Zimmermann, J. (2016) DNAqua-Net: Developing new genetic tools for bioassessment and monitoring of aquatic ecosystems in Europe. Research Ideas and Outcomes, 2, e11321.

Leigh, J. W. & Bryant, D. (2015) POPART: Full-feature software for haplotype network construction. Methods in Ecology and Evolution, 6, 1110–1116.

Lendvay, B., Kadereit, J.W., Westberg, E., Cornejo, C., Pedryc, A. & Höhn, M. (2016) Phylogeography of Syringa josikaea (Oleaceae): Early Pleistocene divergence from East Asian relatives and survival in small populations in the Carpathians. Biological Journal of the Linnean Society, 119, 689–703.

Liska, I. (2015) Managing an international river basin towards water quality protection: The Danube Case. The Handbook of Environmental Chemistry, 39, The Danube River Basin. (ed. by Liška, I.), pp. 1–19. Heidelberg: Springer-Verlag, Berlin, Germany.

Málnás, K., Ambrus, A., Müller, Z., Tóth, Á.P. & Kiss, B. (2011) Bridges as optical barriers and population disruptors for the mayfly Palingenia longicauda: An overlooked threat to freshwater biodiversity? Journal of Insect Conservation, 15, 823–832.

Málnás, K., Polyák, L., Prill, É., Hegedüs, R., Kriska, G., Dévai, G., Horváth, G. & Lengyel, S. (2016) Re-appearance of Palingenia longicauda (Olivier, 1791) (Ephemeroptera, Palingeniidae) on the Hungarian Danube section – range recovery of the species at the Rába-district. Folia Historico-Naturalia Musei Matraensis, 40, 21–25.

Marjoram, P. & Donnelly, P. (1994) Pairwise comparisons of mitochondrial DNA sequences in subdivided populations and implications for early human evolution. Genetics, 136, 673–683.

Marković, V., Kraĉun-Kolarević, M., Kolarević, S., Tubić, B., Ilić, M., Nikolić, V. & Paunović, M. (2017) A first record of Ephoron virgo (Olivier, 1791) (Ephemeroptera: Polymitarcyidae) from the Sava River, with notes on its ecological preferences and rarity of findings in the region. Ecologica Montenegrina, 13, 80–85.

Martynov, A.V. (2018) New records of some rare mayflies (Insecta, Ephemeroptera) from Ukraine. Ecologica Montenegrina, 16, 48–57.

Monaghan, M.T., Inward, D.J., Hunt, T. & Vogler, A.P. (2007) A molecular phylogenetic analysis of the Scarabaeinae (dung beetles). Molecular Phylogenetics and Evolution, 45, 674–692.

Munjiu, O. (2018) Distribution of endangered mayfly Palingenia longicauda (Olivier, 1791) (Ephemeroptera, Palingeniidae) on the territory of the Republic of Moldova, Lauterbornia, 84, 39–51.

Panin, A.V., Astakhov, V.I., Lotsari, E., Komatsu, G., Lang, J. & Winsemann, J. (2020) Middle and Late Quaternary glacial lake-outburst floods, drainage diversions and reorganization of fluvial systems in northwestern Eurasia, Earth-Science Reviews, 201, 103069.

Pauls, S. U., Lumbsch, H. T. & Haase, P. (2006) Phylogeography of the montane caddisfly Drusus discolor: evidence for multiple refugia and periglacial survival. Molecular Ecology, 15, 2153–2169.

Pavel, A.B., Menabit, S., Skolka, M., Lupascu, N., Pop, I.C., Opreanu, G., Stanescu, I. & Scrieciu, A. (2019) New data regarding the presence of two insect larvae species – Gomphus (Stylurus) flavipes (Odonata) and Palingenia longicauda (Ephemeroptera) – in the lower sector of the danube river. Geo-Eco-Marina, 25, 253–264.

Pavlova, A., Beheregaray, L.B., Coleman, R., Gilligan, D., Harrisson, K.A., Ingram, B.A., Kearns, J., Lamb, A.M., Lintermans, M., Lyon, J., Nguyen, T.T.T., Sasaki, M., Tonkin, Z., Yen, J.D.L. & Sunnucks, P. (2017) Severe consequences of habitat fragmentation on genetic diversity of an endangered Australian freshwater fish□: A call for assisted gene flow. Evolutionary Applications, 10, 531–550.

Petit, R.J., Aguinagalde, I., De Beaulieu, J.L., Bittkau, C., Brewer, S., Cheddadi, R., Ennos, R., Fineschi, S., Grivet, D., Lascoux, M., Mohanty, A., Müller-Starck, G., Demesure-Musch, B., Palmé, A., Martín, J.P., Rendell, S. & Vendramin, G.G.. (2003) Glacial refugia: Hotspots but not melting pots of genetic diversity. Science, 300, 1563–1565.

Raymond, M. & Rousset, F. (1995) An exact test for population differentiation. Evolution, 49, 1280–1283.

Rozas, J., Ferrer-Mata, A., Sanchez-DelBarrio, J.C., Guirao-Rico, S., Librado, P., Ramos-Onsins, S.E. & Sanchez-Gracia, A. (2017) DnaSP 6: DNA sequence polymorphism analysis of large data sets. Molecular Biology and Evolution, 34, 3299–3302.

Russev, B. (1987) Ecology, life history and distribution of Palingenia longicauda (Olivier) (Ephemeroptera). Tijdschrift voor entomologie, 130, 109–127.

Schmitt, T. & Varga, Z. (2012) Extra-Mediterranean refugia: The rule and not the exception?. Frontiers in Zoology, 9, 22.

Schwarz, U. (2015) Hydromorphology of the Danube. The Handbook of Environmental Chemistry, 39, The Danube River Basin. (ed. by Liška, I.), pp. 469–479. Heidelberg: Springer-Verlag, Berlin, Germany.

Simons, C., Frati, F., Beckenbach, A. & Crespi, B. (1994) Evolution, Weighting, and Phylogenetic Utility of Mitochondrial Gene Sequences and a Compilation of Conserved Polymerase Chain Reaction Primers. Annals of the Entomological Society of America, 87, 651–701.

Soldán, T., Godunko, R.J., Zahrádková, S. & Sroka, P. (2009) Palingenia longicauda (OLIVIER, 1791)(Ephemeroptera, Palingeniidae): Do refugia in the Danube basin still work?. Communications and Abstracts, SIEEC 21, 81–84.

Stoica, C., Stanescu, E., Paun, I., Banciu, A., Gheorghe, S., Lucaciu, I., Vasile, G.G. & Nita-Lazar, M. (2019) Danube Delta: monitoring and ecological status. A link between the past and the future. Romanian Journal of Ecology & Environmental Chemistry, 1(1), 72–82.

Sworobowicz, L., Mamos, T., Grabowski, M. & Wysocka, A. (2020) Lasting through the ice age: The role of the proglacial refugia in the maintenance of genetic diversity, population growth, and high dispersal rate in a widespread freshwater crustacean. Freshwater Biology, 65, 1028–1046.

Tajima, F. (1989) The effect of change in population size on DNA polymorphism. Genetics, 123, 597–601.

Takenaka, M., Tokiwa, T. & Tojo, K. (2019) Concordance between molecular biogeography of Dipteromimus tipuliformis and geological history in the local fine scale (Ephemeroptera, Dipteromimidae). Molecular Phylogenetics and Evolution. Elsevier, 139, 106547.

Tittizer, T., Fey, D., Sommerhäuser, M., Málnás, K. & Andrikovics, S. (2008) Versuche zur Wiederansiedlung der Eintagsfliegenart Palingenia longicauda (Olivier) in der Lippe. Lauterbornia, 63, 57–75.

Varga, Z. (2010) Extra-Mediterranean refugia, post-glacial vegetation history and area dynamics in Eastern Central Europe. Relict Species: Phylogeography and Conservation Biology (ed. by Habel, J.C. & Assmann, T.), pp. 57–117. Heidelberg: Springer-Verlag, Berlin, Germany

Vartolomei, F. (2012) Integrated measurements for biodiversity conservation in lower Prut Basin. Biodiversity Conservation and Utilization in a Diverse World (ed. by Lameed, G.A.). IntechOpen, London, UK.

Vidinova, Y. & Russev, B. (1997) Distribution and ecology of the representatives of some Ephemeropteran families in Bulgaria. Ephemeroptera & Plecoptera Biology-Ecology-Systematics (ed. by Landolt, P. & Sartori, M.), pp. 139–146. MTL, Fribourg, Switzerland.

Vörös, J., Mikulíček, P., Major, Á., Recuero, E. & Arntzen, J.W. (2016) Phylogeographic analysis reveals northerly refugia for the riverine amphibian Triturus dobrogicus (Caudata: Salamandridae). Biological Journal of the Linnean Society, 119, 974–991.

Voulvoulis, N., Arpon, K. D. & Giakoumis, T. (2017) ‘The EU Water Framework Directive: From great expectations to problems with implementation. Science of the Total Environment, 575, 358–366.

Wattier, R., Mamos, T., Copilaş-Ciocianu, D., Jelić, M., Ollivier, A., Chaumot, A., Danger, M., Felten, V., Piscart, C., Žganec, K., Rewicz, T., Wysocka, A., Rigaud, T. & Grabowski, M. (2020) Continental-scale patterns of hyper-cryptic diversity within the freshwater model taxon Gammarus fossarum (Crustacea, Amphipoda). Scientific Reports. 10, 16536.

Werth, S., Schödl, M. & Scheidegger Christoph (2014) Dams and canyons disrupt gene flow among populations of a threatened riparian plant. Freshwater Biology, 59, 2502– 2515.

Witkowski, Z. J., Król, W. & Solarz, W. (2003) Carpathian List of Endangered Species. WWF and Institute of Nature Conservation, Polish Academy of Science, Vienna-Krakow.

WWF (2020) Living planet report 2020 □ Bending the curve of biodiversity loss. WWF, Gland, Switzerland.

