## Supplementary material for "Cryptic survival and an unexpected recovery of the long-tailed mayfly *Palingenia longicauda* (Olivier, 1791) (Ephemeroptera: Palingeniidae) in Southeastern Europe": Figure S1

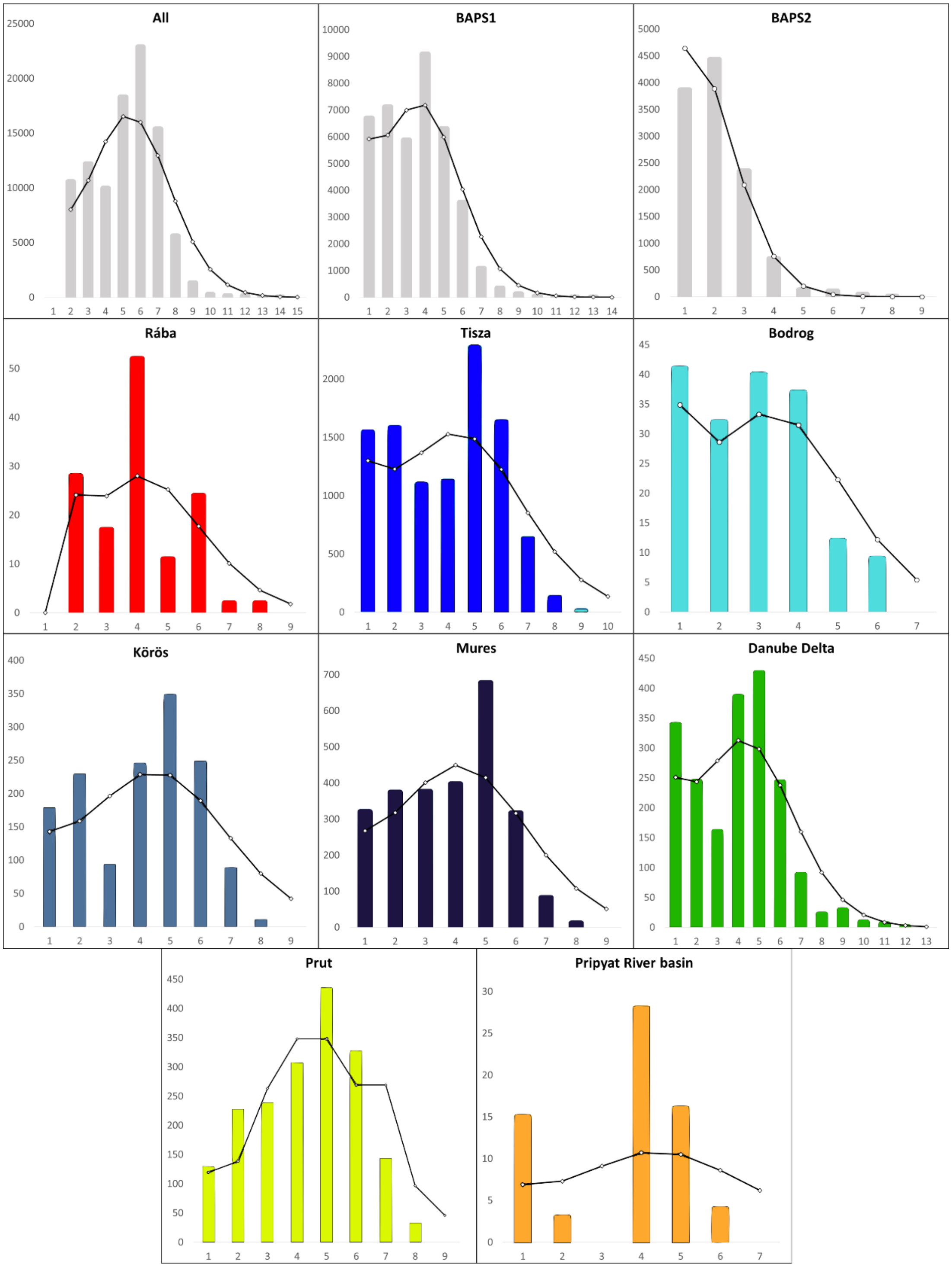


Figure S1. Mismatch distribution histograms, for the whole dataset (All), the two groups identified by the BAPS (BAPS1 and BABS2), and for populations from each river. Bars indicate the observed values and black lines show the expected distribution under the sudden expansion model.
